# Interspecies DNA acquisition by a naturally competent *Acinetobacter baumannii* strain

**DOI:** 10.1101/308833

**Authors:** German M. Traglia, Kori Place, Cristian Dotto, Jennifer S. Fernandez, Camila dos Santos Bahiense, Alfonso Soler-Bistue, Andres Iriarte, Marcelo E. Tolmasky, Robert A Bonomo, Roberto G. Melano, María Soledad Ramírez

**Author notes:** These authors contributed equally to this work (German M. Traglia, Kori Place). Corresponding author. Mailing address María Soledad Ramírez, PhD. Assistant Professor Dept. Biological Science California State University Fullerton 800 N State College Blvd Fullerton, CA 92831.

## Abstract

*Acinetobacter baumannii* is a human pathogen that frequently acquires antibiotic resistance genes leading to the emergence of multi-drug-resistant (MDR) strains. To investigate the role of transformation in the acquisition of resistance determinants by this species, the susceptible strain A118 was exposed to genomic DNA of carbapenem-resistant *Klebsiella pneumoniae* (CRKp). Resistant transformants were obtained and an increase in the resistance level to all β-lactam antibiotics was observed. Whole genome analysis of transformant clones demonstrated the acquisition of CRKp DNA. The most frequently acquired genes correspond to mobile elements, antibiotic resistance genes, and operons involved in metabolism. Bioinformatic analyses and *in silico* gene flow prediction strengthen our findings, showing that a continuing exchange of genetic material between *A. baumannii* and *K. pneumoniae* occurs when they share the same niche. Our results reinforce the idea that natural transformation may play a key role in the increasing emergence of *A. baumannii* MDR.

**IMPORTANCE:** Since the characterization of antibiotic resistance in the late ‘50s, antibiotic resistance propagation was classically associated with horizontal gene transfer (HGT) mediated by plasmids bearing multiple resistance genes. Here we show that, at least in the human pathogen *A. baumannii*, transformation also plays a major role in the acquisition of antibiotic resistance determinants. This study unravels that at least for certain pathogens the propagation of resistance genes occurs by alternative HGT mechanisms which in the past have been unappreciated.

---

During the last few years, the number of antibiotic resistant bacteria prevalent in clinical settings has risen, alarming both scientists and government agencies (1–3). Among these dangerous pathogens, the “ESKAPE” group of bacteria is of profound concern, as these microorganisms cause the majority of hospital infections and effectively “escape” available antimicrobial treatments (4, 5). In this group, “A” represents *Acinetobacter baumannii*, a pathogen that is associated with severe multi-drug-resistant (MDR) infections with attributable mortality levels as high as 60% (through community-acquired pneumonia) and 43.4% (through bloodstream infections) (6–8).

Extreme genome plasticity combined with mechanisms of horizontal gene transfer (HGT)—e.g., conjugation, transformation—have played a key role in the evolution of *A. baumannii*, its adaptability to unfavorable conditions and the acquisition of antibiotic resistance determinants (9, 10). Comparative genomic studies have also shown high variability in *Acinetobacter’s* genome organization, as well as the presence of foreign DNA in their genomes, suggesting exogenous acquisition of genetic traits (11–14). Incorporation of foreign DNA dramatically facilitates the acquisition of antibiotic resistance determinants. These same studies have further identified the presence of the entire DNA uptake machinery in almost all genomes analyzed (13), leading to the assumption that natural transformation could be a common feature in *Acinetobacter* spp. In addition, several studies reported that members of *Enterobacteriaceae* family harbor resistance determinants typically associated to *Acinetobacter* genus, a fact that supports the DNA exchange between these bacteria (15–21). Moreover, recent evidence showed that bacterial predation by *A. baylyi* can cause acquisition of resistance determinants and can be key as a tool for inter-species HGT (22, 23). Our focus of study is the *A. baumannii* clinical strain A118, which was the first *A. baumannii* strain that demonstrated natural competence in its planktonic state (24). It was recently reported that albumin, the main blood protein, and Ca^2+^ are specific inducers of natural competence in *A. baumannii* (2). Other investigations mentioning natural transformation in *A. baumannii* are also known (25, 26). These studies specifically show that DNA uptake occurs only when the isolates were moving on motility medium (26). In sum, genomic studies of clinical isolates as well as experimental natural transformation assays showed that this bacterium can acquire foreign DNA relatively easy.

Hypothesizing that natural transformation plays a role in the acquisition of resistance determinants by *A. baumannii*, we have transformed the *A. baumannii* A118 strain with genomic DNA (gDNA) of two carbapenem-resistant *Klebsiella pneumoniae* (CRKp) clinical isolates (VA360 and Kb18), sequenced the complete genome of selected transformant isolates (A118::VA360 and A118::Kb18) and carried out genomic studies to reveal the acquisition of foreign gDNA. gDNA from *K. pneumoniae* was acquired, including mobile elements, resistance determinants, and genes involved in metabolism. Both bioinformatic analysis and an *in-silico* genome-wide analysis, identified horizontally-derived genes present in both species showing the occurrence of exchange of genetic material between *A. baumannii* and *K. pneumoniae.* Besides, we also observed that *A. baumannii* is capable of acquiring DNA from distant species, such as *Providencia rettgeri* and *Staphylococcus aureus*. This later observation was also backed by *in-silico* genomic analysis. Together, the present data shows that *A. baumannii* can acquire foreign DNA from different species and that transformation is a mechanism implicated in the increasing frequency of emergence of MDR in this threatening pathogen.

## RESULTS AND DISCUSSION

### Natural transformation of the *A. baumannii* A118 strain with gDNA of CRKp VA360

To confirm the role of natural transformation in the acquisition of resistance determinants in *A. baumannii*, we performed natural transformation assays using the *A. baumannii* A118 strain as acceptor as previously described (24). We used 200 ng of gDNA of VA360 strain, whose complete genome is available (NZ_ANGI00000000.2), as donor DNA. Previous reports have demonstrated that *K. pneumoniae* VA360 is multidrug resistant (MDR) harboring five plasmids and four class A β-lactamases including *bla*_TEM-1_, *bla* _KPC-2_, *bla*_SHV-11_, *bla*_SHV-12_ (27). The acquisition of DNA could lead to the emergence of a resistant phenotype in *A. baumannii* A118. Importantly, transformant isolates were recovered at a frequency of 8.38 × 10^−7^ (SD± 5.64 × 10^−7^) in LB agar plates containing 1 μg/ml of meropenem (MEM). Disk diffusion was used as the screening method to explore additional changes in the resistance phenotype, one transformant colony (A118::VA360) was selected for further studies. Antimicrobial susceptibility profiles showed that susceptibility was modified for the β-lactam antibiotics tested, including MEM and imipenem (IPM) (Table 1). The selected transformant showed increased resistance to MEM and imipenem (IPM). Interestingly, an increase in the inhibition zone was observed for other antibiotics such as SXT and NAL (Table 1). Changes in antibiotic susceptibility were confirmed by MIC determination. As shown in Table 2, MEM MIC increased 100 times from 0.125 μg/ml in A118 to16 μg/ml in A118::VA360. IPM MIC the increased 5 times from 0. 25 μg/ml to 1.5 μg/ml (Table 2). To determine which DNA acquisition events led to changes in susceptibility in A118::VA360, its whole genome was sequenced.

**Table 1.**
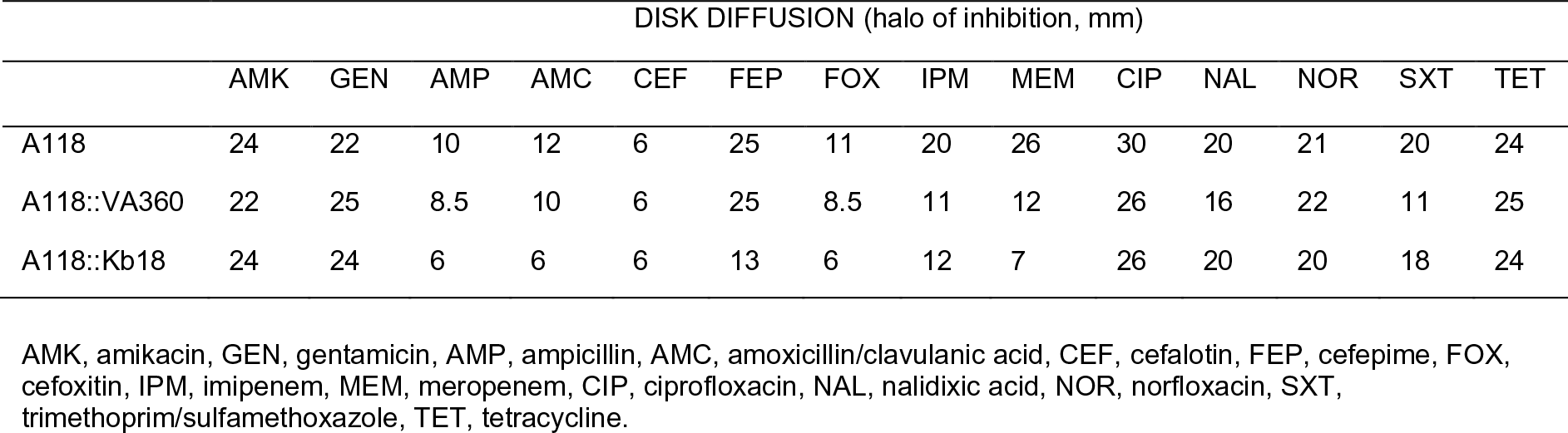
Antibiotic susceptibility test (disk diffusion) of A118 and A118 transformed cells with DNA of *K. pneumoniae* strain VA360 and Kp18 (A118::VA360 and A118::Kb18)

**Table 2.**
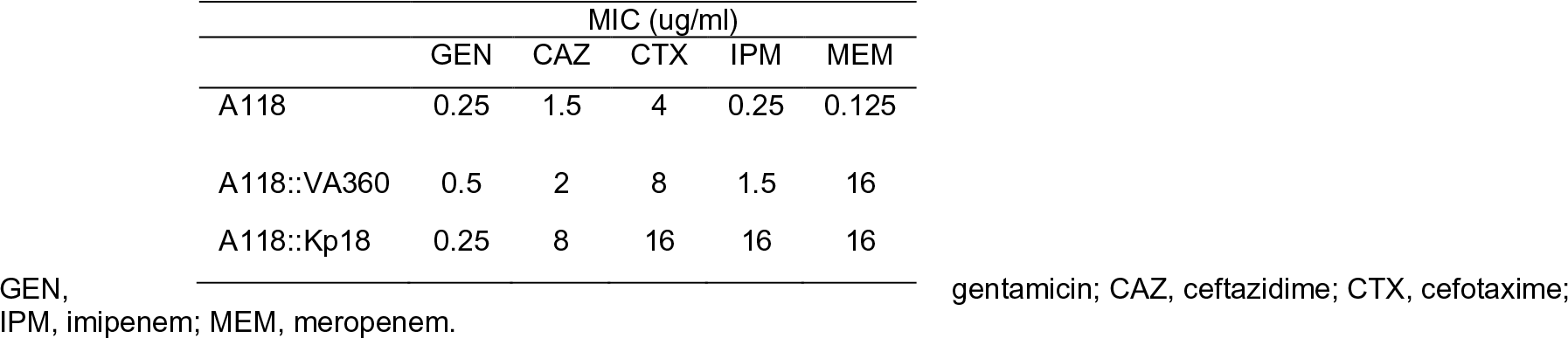
Antibiotic susceptibility test of A118 and A118 transformed with DNA of *K. pneumoniae* strains VA360 and Kb18.

### Genomic analysis and distinctive features of A118::VA360 transformant

To characterize in depth the genotypic changes underwent by A118, the complete genome sequence of the one selected transformant as well as A118 wild type (WT) strain were obtained to perform genomic comparison. The general features of the draft genomes are summarized in Table 3.

**Table 3.** General features of A118, A118::VA60 and A118::Kb18.

| Strain | A118 | A118::VA360 | A118::Kb18 |
| --- | --- | --- | --- |
| Size (bp<sup>a</sup>) | 3,853,242 | 3,885,049 | 4,058,089 |
| G+C contents (%) | 38.69 | 38.81 | 38.65 |
| ORFs<sup>b</sup> | 3,589 | 3,819 | 4,058 |
| tRNA | 60 | 49 | 72 |

<sup>a</sup>bp, base pairs.
<sup>b</sup>ORFs, Open reading frames.

Forty-seven DNA fragments, represented by 6,205 bp, that are not present in A118 genome, were found in A118::VA360 genome. These DNA fragments varied in size with an average size of 709 bp, maximum fragment size of 4,367 bp and minimum fragment size of 75 bp (Fig. 1).

**Fig. 1.**
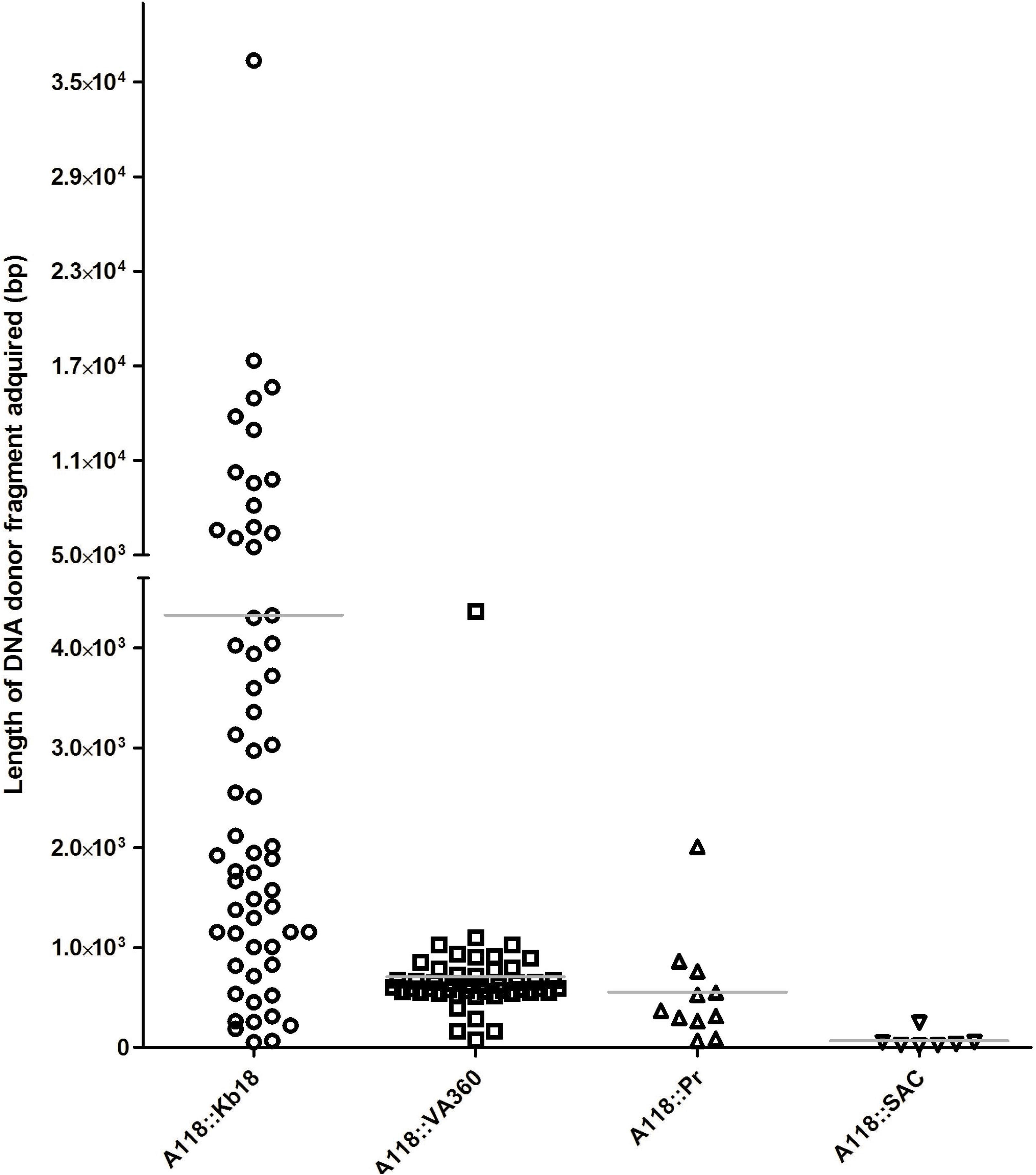
Dot plot representing the distribution of DNA-acquired fragment lengths in *A. baumannii* transformant isolates (A118::Kb18, A118::VA360, A118::Pr and A118::SAC). The arithmetic mean was indicated by a line. Circles represent Kb18 acquired-DNA, squares represent VA360 acquired-DNA, triangles represent Pr acquired-DNA, and inverted-triangle represents SAC acquired-DNA.

We next compared the 47 DNA fragments imported into the genome of A118 to 5,432 *A. baumannii* genome sequences from GenBank (excluding A118 genome). Twelve out of the 47 imported DNA fragments have been previously reported in *A. baumannii* genomes. Acquired fragments of DNA where we confirmed an insertion site in A118 genome corresponded to hypothetical proteins and intergenic regions from *K. pneumoniae* VA360 strain (Table S1). Furthermore, mobile genetic elements known to be present in *K. pneumoniae* VA360 were found among the imported DNA fragments identified in the A118::VA360 draft genome sequence. The genetic elements found in A118::VA360 were: IS*Aba14*, IS*Kpn26*, IS*CR1* (IS*91* family), IS*26*, IS*1R* and the Tn*3* transposon. In addition, the common 3’ conserved region of the class 1 integron with *sul1* gene conferring sulfonamide resistance and *qacEΔ1* gene conferring ammonium quaternary resistance were present. Another resistance gene that we identified was *aac(6’)-lb-cr*, which confers resistance to aminoglycoside and reduced susceptibility to quinolones. In addition, six genes encoding efflux pumps previously reported in *K. pneumoniae* strains were found in the A118::VA360 transformant.

Also of importance, we identified eight genes related with metabolic pathways in A118::VA360 (Table S1). Among them, we found the *hpaX* gene that encodes 4-hydroxyphenylacetate permease, an enzyme associated with the *hpa* gene cluster responsible of 4-hydroxyphenylacetate (4-HPA) metabolism. Arcos et al., reported that the *hpa* cluster has been found in several microbes with different genetic organization, including two *hpa* clusters in *K. pneumoniae*, one containing only the *hpaR* gene, and the second containing *hpaG1G2EDFHIXABC* genes (28). 4-HPA is a small, aromatic molecule secreted in human saliva (29), found to induce NadA expression in *Neisseria meningitidis* and subsequently promote bacterial adhesion in the oropharynx niche (30). Although we identified the *hpaX* in A118::VA360 other genes of *hpa* cluster were not found. While it is possible that *hpaX* alone could enhance virulence, its function in the absence of the *hpa* gene cluster is unclear.

Additionally, we identified the *cutA* gene that encodes for a dihydroorotate dehydrogenase, which has an enzymatic function associated with copper tolerance and osmotic stress (31, 32), but also related to plasmid and chromosomal replication (31, 33). This can explain the potential of *cutA* to be transferred between different bacterial species. Furthermore, *fieF* (also called *cepA* or *yiiP)*, which encodes for a ferrous iron efflux system, and it is also reported as an iron and zinc detoxification efflux system (34, 35) was identified in A118::VA360. Iron chelation is utilized by the innate immune-system to limit the iron available to invading microbes. Subsequently, bacteria must circumvent iron limitations, as FieF is an iron transporter, could be considered a virulence factor found in several pathogenic bacteria (36, 37). Other studies have also linked *fieF* to disinfectant resistance phenotype in *K. pneumoniae* (38–40). Consequently, the acquisition of FieF in *A. baumannii* could enhance survival and persistence in the clinical setting. Also, we observed the presence of nine fragments of DNA containing genes with unknown function and intergenic regions.

Recent evidence has suggested that bacterial predation by *A. baylyi* can facilitate the acquisition of resistance determinants and may play a key role in interspecies HGT (22, 23). To test the role of bacterial predation in *A. baumannii* the ability of A118 to lyse *K. pneumoniae*, as well as *E. coli* was assessed by performing killing assays. A118 was used as predator and VA360 or *E. coli* MG1655-Rif as preys as previously described (41, 42). We clearly observed that A118 kills VA360 (Fig. S1A and B). Our results showed A118 killing activity against both species tested, showing a dramatic predation towards *E. coli* (Fig. S1B). Future experiments to observe *in situ* DNA acquisition will be performed.

### Natural transformation assays using genomic DNA from CRKp Kb18 harboring *bla*_KPC_ and *bla*_OXA-23_

Since we observed the existence of interchange between *A. baumannii* and *K. pneumoniae*, and because our previous transformant A118::VA360 developed increase resistance to carbepenems not acquiring carbapenemase activity, we decided to test if there is a preferential selectivity to acquire some carbepenemases over others. For these reasons, and to further test our hypothesis, natural transformation assays using A118 strain by adding gDNA of another CRKp strain (Kb18) known to harbor two carbepenemases, *bla*_KPC_ and *bla*_OXA-23_, were performed.

Transformation assays resulted in a frequency 7.17 × 10^−7^ (SD± 1.89 × 10^−7^) CFU/ml in LB agar plates containing 1 μg/ml of MEM. As stated previously, after an initial susceptibility screening, one transformant colony (A118::Kb18) was selected for further studies. Like A118::VA360, increased levels of resistance to all β-lactams were observed for A118::Kb18 (Table 1 and 2)

The complete genome sequence of A118::Kb18 was obtained to perform genomic comparison and confirm the presence of resistance determinants and acquired DNA. The general features of the A118::Kb18 draft genome are summarized and compare with A118 and A118::VA360 genomes in Table 3.

Sixty-two new DNA fragments were acquired by A118::Kb18 strain. The average size of DNA fragments acquired by natural transformation was 4,331 bp with a maximum fragment size of 36,369 bp and minimum fragment size of 1,042 bp (Fig. 1).

Focusing our analysis on resistance determinants and mobile elements, we observed that A118::Kb18 had acquired six genes coding for resistance to β-lactams (*bla*_TEM-1_ and *bla*_OXA-23_), streptomycin, spectinomycin (*strA, strB, aadA1, sat2*) and trimethoprim (*dfrA1*). The acquisition of *bla*_OXA-23_ but not of *bla*_KPC_ suggests that *A. baumannii* may have a preferential affinity to acquire and maintain certain carbapenemases over other. The observed results agree with the observation that *bla*_OXA-like_ enzymes are frequently found in this species and only a few reports of *bla_KPC_ A. baumannii* positive isolates were reported in the literature (41, 42). The *intI2* gene together with the typical gene cassette array of the class 2 integron In2-7, *dfrA1-sat2-aadA1-orfX-ybfA-ybfB-ybgA*, was found. Moreover, the complete transposition module (*tnsE, tnsD, tnsC, tnsB* and *tnsA*) of the Tn*7* transposon was also identified. In addition of the Tn*7* transposon, A118 acquired other mobile elements was observed. Eight IS and three transposons where detected among the 62 DNA imported DNA fragments (Table S1).

Similarly, *bla*_TEM-1_ was found into the typical structure of Tn*3* transposon. The IS*Aba125*, which was absent in A118 genome prior exposition to Kb18 gDNA, was also found in the transformant isolate. The presence of IS*Aba125* and *bla*_OXA-23_ in the gDNA source (Kb18) and in the selected transformant cell, called our attention and highlight the importance of search for these genes. This led us to perform a retrospective surveillance to identify the presence of *bla*_OXA-23_, IS*Aba125*, and IS*Aba1* in a collection of twenty-two CRKp isolates. Three CRKp strains (Kp16, Kp8, Kp21) were positive for bla_OXA-23_ by PCR and Sanger sequencing. Moreover, bioinformatics analyses searching for shreds of evidence of the presence of *bla*_OXA-23_, *bla*_OXA-24_, *bla*_OXA-51_, *bla*_OXA-58_, IS*Aba125*, and IS*Aba1* were performed. With the exception of *bla*_OXA-58_, the presence of all these genes and IS was observed in K. pneumoniae sequences deposited in the GenBank database (Table S2 and S3).

Correspondingly, the presence of *bla*_OXA-23_, *bla*_OXA-24_, *bla*_OXA-51_, and *bla*_OXA-58_, was also investigated in Enterobacteriaceae genomes deposited in the GenBank database. In addition to *K. pneumoniae*, we found that only one sequence of E. coli and two sequences of *Proteus mirabilis* possessed *bla*_OXA-23_ (Table S3). The *E. coli* sequence (accession number KJ716226) corresponds to a non-self-conjugative plasmid of 50-Kb (16) where bla_OXA-23_ was associated with the *IS1*. The *E. coli* strain (*E. coli* 521) was recovered from a urine sample of an elderly female and possessed other ß-lactamases (16). Moreover, Paul et al. described the presence of fourteen E. coli isolates harboring *bla*_OXA-23_ obtained from patients hospitalized in the intensive care unit (18). In P. mirabilis strain CFO239, bla_OXA-23_ was inserted into the chromosome (43), while in the extended-spectrum β-lactamase, ESBL, P. mirabilis strain 4969, the authors suggested a plasmid location of bla_OXA-23_ (21). bla_OXA-58_ was also found in three P. mirabilis, two of them harbored in plasmids (15), while the other was associated with a *bla_AmpC-like_* gene both into an integrated prophage (19). Leski et al also reported the presence of *bla*_OXA-58_ in members of the Enterobacteriaceae family: five Enterobacter spp., one *E. coli* (SL-5) and three K. pneumoniae isolates (20). Six of these strains also harbored bla_OX_A_51-like_ genes apart of *bla*_OXA-58_ (20). The *bla*_OXA-58_ sequence found in *Enterobacter* sp. strain SL1 is the only sequence submitted to GenBank (KC004135).

We also searched for IS*Aba1* and IS*Aba125* genes among *Enterobacteriaceae* genomes successfully finding both elements (Table S3). In our search we found IS*Aba1* in *K. pneumoniae, E. coli, Salmonella enterica* and *Shigella flexneri*. IS*Aba125* was more frequently observed in more diverse bacterial species: we found it in *E. coli, K. pneumoniae, Klebsiella* spp., *S. enterica, Enterobacter* spp., *Citrobacter freundii, Cronobacter sakazakii*, and *Raoultella planticola* (Table S3). Analyzing the genome of 1,777 ESBLs positive *K. pneumoniae* strains, Long et al. observed the presence of OXA genes (OXA-23, OXA-24, OXA-48, and OXA-83) in 23 strains (44). Five contained OXA-23, and five were positive for OXA-24. Collectively, our results and the present lines of evidence highlight the importance of investigating genes that are frequently reported in certain species in others that share the same niche. Furthermore, a genetic platform that contains several IS, such as IS*CR2*, ΔIS*CR2* and IS *1006* and the genes *strA, strB* and *floR* was also acquired by A118::Kb18. Different genetic structures sharing some of the elements found in A118::Kb18 were also described previously in *K. pneumoniae* (Fig. 2).

**Fig. 2.**
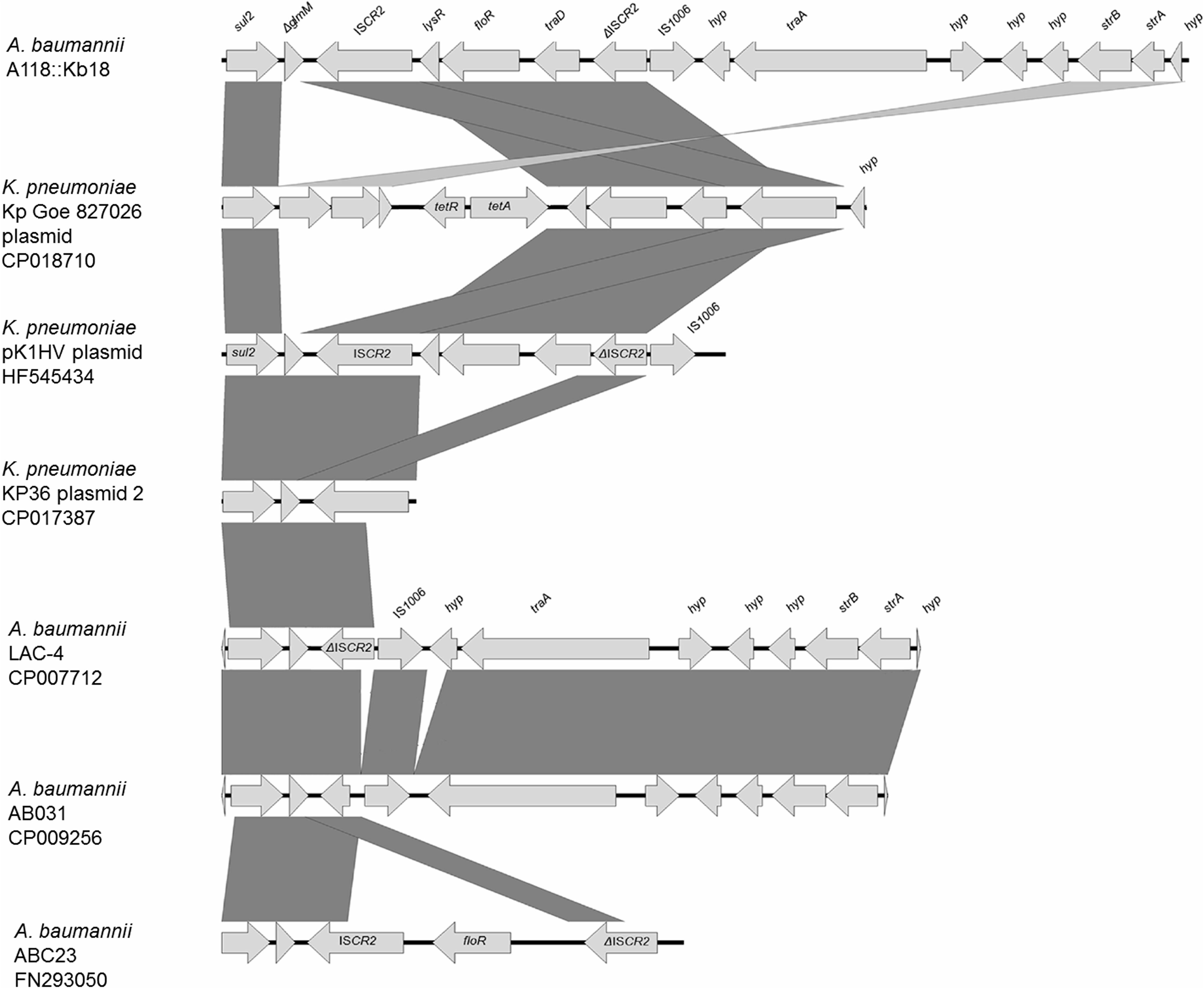
Genetic structure and comparison of IS*CR2* and its associated antibiotic resistance genes. The genetic structure of A118::Kb18 was compared with similar genetic structures from *K. pneumoniae*. The graphic representation was made using the EasyFig version 2.2.0 software.

The presence of IS*CR2* was also examined in the collection of our twenty-two CRKp, and six isolates were positive.

Considering all the exposed results, and the evidence of genes commonly found in *A. baumannii* in our CRKp isolates and *K. pneumoniae* sequences, we searched for the presence of gene sequences most generally reported in *Enterobacteriaceae* in *A. baumannii* genomes. The presence of *bla*_KPC_, *bla*_TEM-1_, *bla*_SHV-2_, *bla*_CTX-M-9_, *bla*_CTX-M-2_, *bla*_CTX-M-1_, ISCR1 and ISEcp1 was investigated, and we found all of them in at least one *A. baumannii* genome (Fig. 3 and Table S3-4). We observed some distinct features for some of the genes, e.g., the *bla*_CTX-M-9_ gave us only 48% of coverage. However, it has 94-99% identity in the six obtained matches. All the isolates harboring this bla_CTX-M_ were recovered in China (Fig. 3 and Table S3). Another feature that we have observed was the presence of *bla*_KPC_ mainly in *A. baumannii* strains that were isolated in Brazil; only three of the 24 findings correspond to USA isolates.

**Fig. 3.**
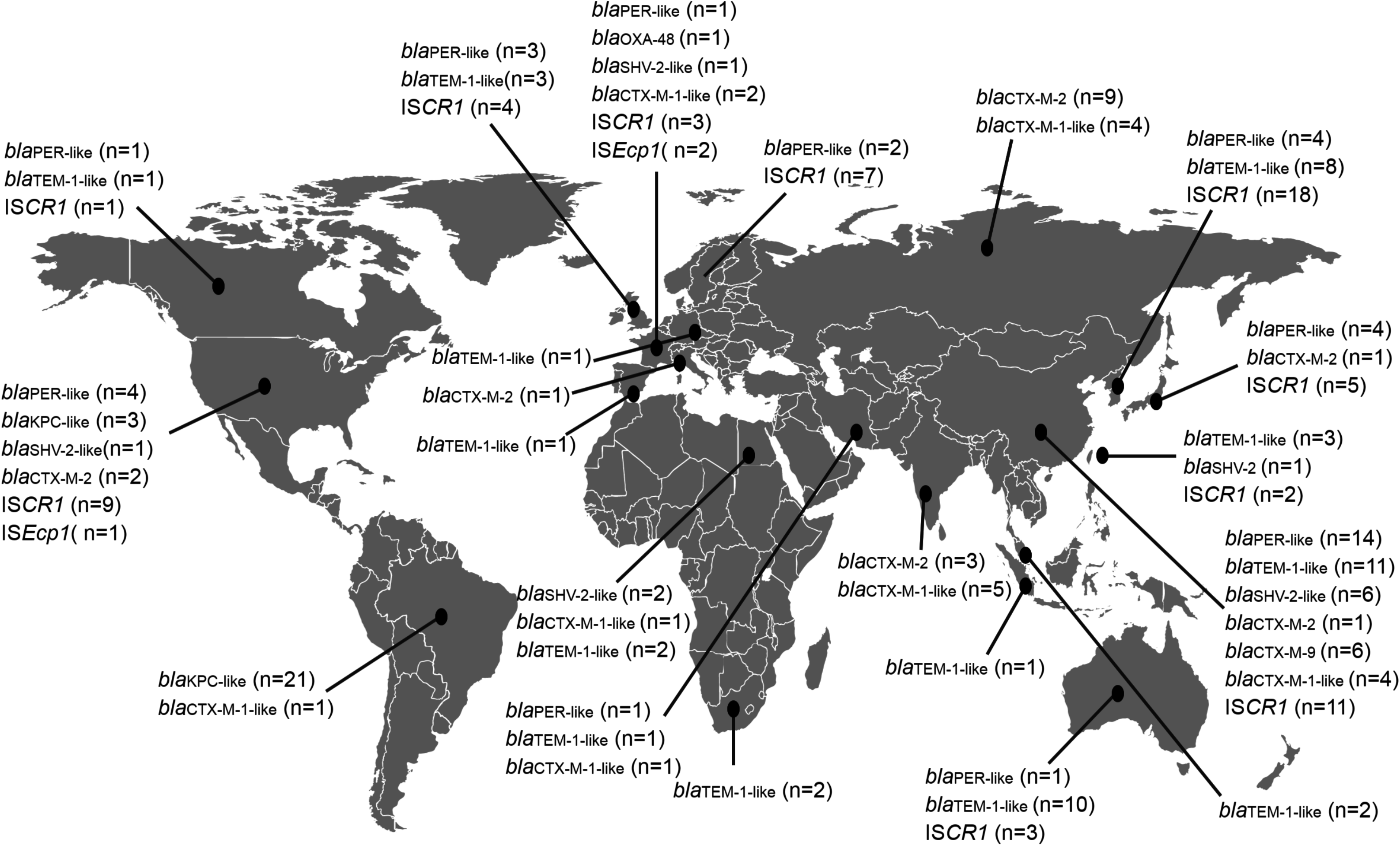
Global distribution of gene sequences most generally reported in *Enterobacteriaceae* in *A. baumannii* genomes.

In agreement with our observations, Ramirez et al, described the ability of A. baumannii strain A118 to gain and maintain a plasmid harboring *bla*_CTX-M-2_ gene, which was previously found in several P. mirabilis isolates (45). Although there are few reports describing the presence of *bla*_CTX-M-2_ in A. baumannii isolates this work shows the ability of A. baumannii to gain and maintain plasmids carrying genes commonly found in Enterobacteriaceae.

In addition, by PCR and using 23 A. baumannii strains from our collection, we searched for the most commonly described insertion sequence (ISKpn1) among *K. pneumoniae* isolates. No positive results were obtained in the tested *A. baumannii* strains.

To identify other genetic elements related to HGT, we carried out analysis of putative phage sequences in A118 and A118::Kb18. Out of 6 prophages detected in A118::Kb18, four of them were acquired when A118 was transformed with Kb18 gDNA. Interestingly, we identified the insertion of putative prophage of 36,369 bp localized in a gene that codes for a hypothetical protein upstream to *tonB* gene cluster. The insertion of putative phage upstream *tonB* in A118:Kb18 genome should modify the genetic expression of it. In Gram-negative bacteria, the ExbB-ExbD-TonB system is responsible to provide the energy to transport host iron-carrier and iron-siderophore complexes into the periplasm once these complexes are bound to cognate TonB-dependent outer membrane receptors. Zimbler et al 2013 described three copy of an active transcriptionally *tonB* gene into different genetics structure (46).

The presence of a putative group II intron, not detected in A118 genome, was found into A118::Kb18 genome. Group II introns are catalytically active RNA and mobile genetic elements present both in eukaryotic and prokaryotic cells. These introns were observed in different *A. baumannii* and *K. pneumoniae* strains and were present within the variable region of class I integrons (47, 48). The group II intron found in A118::Kb18 was also present in a great number (n=125) of *A. baumannii* genomes deposited in the GenBank database. In the A118::Kb18 strain, this intron was found inserted between genes that codify a hypothetical protein and TonB-dependent siderophore receptor. This group II intron was also present in three *K. pneumoniae* genomes linked with class I integron genetic structures. Moreover, when we compared our sequence with the ones described in the GenBank, the same genetic structure was only found in *A. baumannii* 11510 (CP018861), AF-401 (CP018254), AbH12O-A2 (CP009534) and AB030 (CP009257) genomes.

The analysis of the A118::Kb18 genome also showed a great number of DNA fragments that do not correspond to ISs, transposon or putative prophage sequences or were related to these elements (total sum of 87,374 bp). These DNA fragment that are not related with mobile elements might be inserted into the genome by homologous or illegitimate recombination. The in-depth analysis of the nucleotide sequence of these acquired DNA fragments showed that 46,365 out of 87,374 bp corresponded to genes with unknown functions or sequences that were identified as intergenic regions of *K. pneumoniae* genomes. However, we have also observed some fragments (n=9) containing genes or cluster of genes related to virulence traits or metabolic processes (Table S1). Among the above mentioned fragments with a defined role, a region of 12,946 bp containing the genes related to 4-hydroxyproline uptake and utilization was found. The 4-hydroxyproline is the principal component of collagen. Wong et al. reported that transurethral colonization of uropathogenic *E. coli* into male mice induced persistent prostatic inflammation followed by a significant increase in collagen deposition and hydroxyproline content (49). In order to identify if this cluster is expressed in A118:Kb18, the acquisition and utilization of 4- hydroxyproline of A118::Kb18 compared with the isogenic strain was performed. We observed that A118::Kb18 utilized 4-hydroxyproline more efficiently in comparison to A118 WT strain (Fig. 4). Therefore, the acquisition of this gene cluster will contribute in the virulence on A118 transformant clone.

**Fig. 4.**
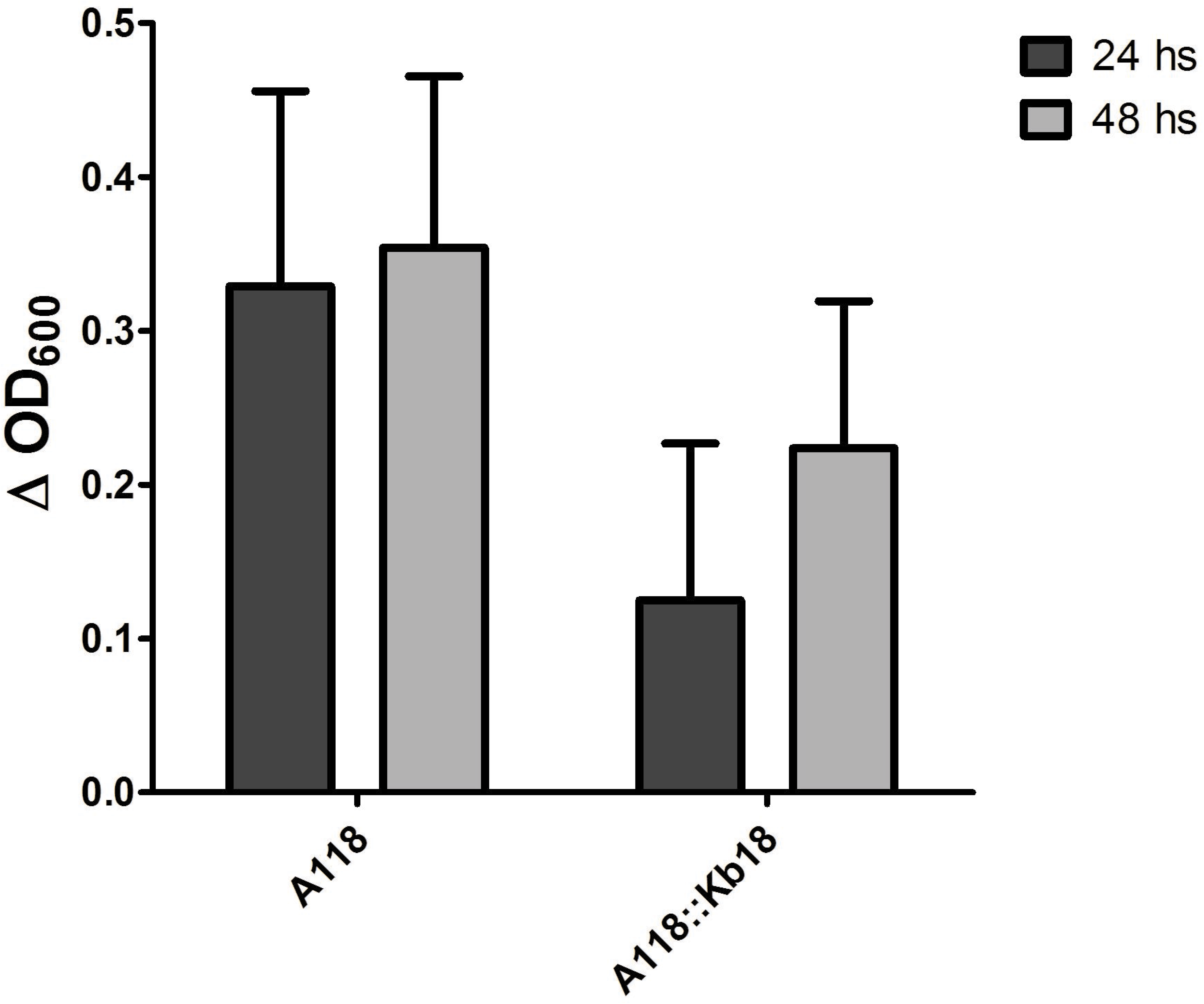
Utilization of 4-hydroxyproline of A118::Kb18 compared with the A118 WT strain.

Since transformation with foreign DNA could cause genomic changes in A118 that can result in different phenotypes, total proteins and exo-proteins of A118 and A118:Kb18 we obtained. Differences were not identified in the total protein and exo-protein patterns when we compared A118 against A118::Kb18 (Fig. S2). Moreover, the ability to form biofilms on polystyrene surface by these two isolates was also investigated showing no significant changes in relation to the ability to form biofilms by A118 and A118::Kp18 (Fig. S3).

A significant amount of evidence is presented of DNA-uptake by A118 that was obtained from the genomic analysis that has an impact in the resistance profile. Additionally, genes encoding for unknown proteins and not well defined features were also incorporated, increasing the pathogenic potential.

Our results reinforce the idea that natural transformation could be involved in the increasing emergence of AMR in the threatening pathogen, *A. baumannii*.

### Growth *A. baumannii* A118 transformant isolates is not impaired

Growth curves for the A118, A118::VA360, and A118::Kb18 were performed (Fig. S4). Differences in growth rates were not observed, indicating that the massive acquisition of foreign DNA by A118 (e.g. ~0.2 Mb in the case of A118::Kb18) did not affect the growth of the bacterial cells. These results suggest that *A. baumannii* could acquire long fragments of DNA by natural transformation without losing its fitness, which would increase its chances of competing against other microorganisms for the same niche.

### Supporting evidence of genetic exchange between *A. baumannii* and *K. pneumoniae* by HGT

To further support and validate the gene flow and the interplay between *A. baumannii* and *K. pneumoniae*, predictive analysis was performed using 15 and eight complete genomes from the Genbank, respectively (Table S5). We determined HGT-acquired genes by phylogenetic tree reconstruction and reconciliation analysis to predict of HGT events. To make this analysis more stringent we excluded mobile gene elements (such as ISs), integrons, pseudogenes, phage related sequence and intergenic regions.

We observed an average of 14 (11–18) horizontally transferred genes per genome from *K. pneumoniae* to *A. baumannii*. For example, the genome of *A. baumannii* strain ZW85-1 showed the presence of 18 horizontally transferred genes; in comparison, strain AB0057 contained 11 genes that were transferred (Table S3).

Considered individually, our *in-silico* analysis predicted that an average of 11 (9-14) genes with known function (e.g. *rpoN, rhtC, lpxB*) and two (1–4) genes with unknown function would be transferred per genome (Table S3).

We compared the candidate genes predicted by the *in-silico* tree reconciliation analysis to our experimental analysis of A118::VA360 and A118::Kb18 genomes and found similarities between the two (some of the genes transferred into A118::VA360 and A118::Kb18 were also present in our *in-silico* analysis). For example, *hpa* was predicted to be transferred into *A. baumannii* ZW85-1 genome. In agreeance with the tree reconciliation analysis, *hpa* was found in the genome of A118::VA360 transformant. The putative threonine efflux protein (*rhtC*) and the lipid A disaccharide synthetase (*lpxB*) from *K. pneumoniae* was predicted to be transferred into all 15 *A. baumannii* genomes from GenBank. In agreement, *rhtC* and *lpxB* were transferred into the A118::Kb18 transformant. Similarly, the transfer of one putative transcriptional regulator belonging to the MerR family from *K. pneumoniae* was predicted into 12 *A. baumannii* genomes. As expected, MerR was found in the A118::Kb18 transformant.

Next, we assessed the occurrence of reciprocal gene flow between *A. baumannii* to *K. pneumoniae*. Notably, we observed an average of 30 genes (27–34) transferred from *A. baumannii* to *K. pneumoniae*. Among them, an average of 28 genes (25–35) have known function (e.g. *aroB, dapE, dprA, gpml*) and a range of 1 to 4 genes possess unknown function (hypothetical protein) in average per genome (Table S5). Considering our gene flow in-silico analysis, gene transfer is bidirectional albeit, *K. pneumoniae* is more prone to acquire genes from *A. baumannii* than vice-versa.

Our results expose the dynamic and frequent exchange of genetic material between two species of Gamma-proteobacteria that share the same niche. Therefore, exchange of genetic material could be a consequence of the continuous interplay between *A. baumannii* and *K. pneumoniae* in a clinical setting.

### Further evidences of A118 DNA acquisition using other DNA sources

To further study *A. baumannii’s* capability to acquire DNA from other MDR bacteria, we performed transformation assays using *Providencia rettgeri* strain M15758 (harboring *bla*_NDM-1_) and the methicillin-resistant *Staphylococcus aureus* “Cordobes” clone (SAC) strain CD, as DNA sources. Transformation assays, using 10 μg/ml of cefotaxime and 200 μg/ml of ampicillin as antibiotic selection, were performed as previously described, and two selected transformant colonies were sent to whole genome sequencing (A118::Pr and A118::SAC).

Global analysis confirmed the acquisition of foreign DNA from both DNA sources. A variable length of acquired DNA fragments was observed. The average size of DNA fragments acquired by natural transformation in A118::Pr was 556 bp, ranging from 67 bp to 2,011 bp. We observed DNA acquisition from non-coding sequences (n=3) integrated in intergenic regions of A118 genome, as well as the acquisition of several DNA fragments containing genes related with metabolic pathways (acetyl-CoA acetyltransferase) or oxidative stress (alkyl-hydroperixode reductase) (Table S1). Strikingly, we found *pilJ* and *pilK* genes from *Salmonella enterica*, both associated with type IV pilus biogenesis in this species (50). In addition, we found the acquisition of an aminoglycoside resistance gene, *aadB*, preceded by the class 1 integron integrase.

Only four DNA transfer events were observed in the A118::SAC transformant cell. All four events corresponded to non-coding sequences that were found in the intergenic region in the A118 genome (Table S1). The average size of the acquired DNA fragments in A118::SAC was 67 bp with a maximum fragment size of 250 bp and minimum fragment size of 21 bp. These results suggest that a low homology sequence and a great phylogenetic distance between two species plays an important role in DNA acquisition into the *A. baumannii* genome.

As previously performed, tree reconciliation analysis (51) were used to explore the occurrence of HGT events between *A. baumannii* and S. *aureus* using available genomes in Genbank. For this purpose, eight *A. baumannii* genomes and 37 S. *aureus* genomes were used (Table S5).

We observed the presence of one to two horizontally transferred genes per genome from S. *aureus* to *A. baumannii* (Table S5). All the predicted transferred genes obtained by *in-silico* analysis possessed a known function. Among them topoisomerases, ATPases and a gene related with capsular polysaccharide biosynthesis were identified (Table S5).

The occurrence of reciprocal gene flow from *A. baumannii* to *S. aureus* showed an average of two horizontally transferred genes (1-3 genes) from *A. baumannii* to S. *aureus*. Among these genes all of them have a known function, such as topoisomerases, phosphopantothenoylcysteine decarboxylases, ligases, and genes involved in capsular polysaccharide synthesis among the others (Table S5). Accordingly, a few HGT events were observed in the experimental assay and through the in-silico genome-wide analysis, suggesting an infrequent gene flow between *A. baumannii* and *S. aureus*.

We also observed a unique trend in DNA acquisition by *A. baumannii*. This could be explained by a secondary event of homologous recombination after DNA-uptake. A tendency to acquire non-coding DNA fragment than coding sequences prevailed. Nevertheless, this suggests that the non-coding sequences could generate a new target for additional recombination events within the *A. baumannii* population. Although, non-homologous recombination mediated by mobile element or illegitimate recombination occurs in *A. baumannii* fact that was observed in the transformant isolates (A118::Kb18, A118:VA360 and A118:Pr). These results coincide with Dominguez et al. findings that natural transformation in *A. baylyi* ADP1 plays an essential role in the acquisition of mobile genetic elements (52). Also, those results suggested that the DNA from Gram-negative bacteria served as a preferred source of genetic material than DNA from Gram-positive bacteria, that can ultimately contribute in the evolution of *A. baumannii*.

## CONCLUSION

Our results show that *A. baumannii* A118 can successfully acquire and recombine foreign DNA from other species, leading to, among known phenotypic features, a change in the susceptibility antibiotic resistance profile. Even if these results were obtained *in vitro*, the rate of transformation indicates that this mechanism of HGT is a common way for adaptation and evolution of this species. *In silico* analysis support these findings, indicating a “two-direction” genetic flux between gram negative bacteria sharing the same ecological niche. Previouly, bioinformatic analysis and in silico prediction showed that different species of pathogens could share resistance determinant. Here we showed that *A. baumannii* readily acquired resistance determinants, and other virulence traits, from *K. pneumoniae* strains. Thus, *K. pneumoniae’s* virulence properties could be followed by the capture of its genomic material. More work is needed to prove our results point out towards this direction.

Previous reports using the *A. baylyi* strain showed the acquisition of different mobile elements -such as transposon, gene cassettes- from different species (52). In addition, recent studies showed the ability of *A. baylyi* to use the T6SS to lyse and acquire genes from neighboring cells. These studies showed the important and contribution of bacterial predation on cross-species HGT (22, 23). In summary, this study and other previous findings, reinforce the idea that transformation could play an important role in the evolution of *A. baumannii* towards the MDR and that this mechanism could be implicated in the increasing frequency of emergence of MDR in this threatening pathogen.

## MATERIAL AND METHODS

### Bacterial strains and standard molecular biology techniques

The naturally competent carbapenem-susceptible clinical strain *A. baumannii* A118 was used in the transformation assay as the recipient (2, 24). Two carbapenem-resistant *K. pneumoniae* (CRKp) strains (VA360 and Kb18) were used as the gDNA source to transform A118 (27). *K. pneumoniae* VA360 was isolated in 2006 from a tertiary care medical center in Cleveland, OH, USA (27). Previous reports have demonstrated that VA360 is multidrug resistant, harboring four class A β-lactamases including *bla*_TEM-1_, *bla*_KPC-2_, *bla*_SHV-11_, and *bla*_SHV-12_ (27). The other strain, *K. pneumoniae*, strain Kb18, was isolated in 2014 from an intensive care unit in Buenos Aires, Argentina. This strain was multidrug resistant and was positive for *bla*_KPC_ and *bla*_OXA-23_. Total DNA extractions from donor cells (VA360 and Kb18) were carried out using Wizard^®^ Genomic DNA Purification Kit according to manufacturer instructions (Promega, Madison, WI). PCR reactions were carried out to confirm the presence of resistance determinants previously identified in the donor strains. The reactions were performed using the Zymo Taq™ PreMix following manufacturer’s instructions (Irvine, CA, USA). Other DNA sources used for transformation were DNA from *Providencia rettgeri* strain M15758 (*bla*_NDM-1_) and *Staphylococcus aureus* “Cordobes” clone (SAC) (*mecA*) (53, 54).

Moreover, a total of 22 CRKp *K. pneumoniae* and 23 *A. baumannii* strains randomly selected from our collection were used to search for the presence of *bla*_OXA-23_, IS*Aba125*, IS*Aba1* and IS*CR2*; and IS*Kpn1*, respectively. PCR reactions using specific primers were carried using the Zymo Taq™ PreMix according to manufacturer’s instructions (Irvine, CA, USA).

### Natural transformation assays

Standard natural transformation assays were performed as previously described (24). Briefly, 50 μl of late stationary-death phase cultures of *A. baumannii* A118 were transferred to 50 μl of sterile LB with 200 ng of each donor gDNA. These cultures were incubated for 1 hour at 37°C and then plated on LB agar with 1 μg/ml of meropenem (MEM), 10 μg/ml of cefotaxime or 200 μg/ml of ampicillin, with the goal of selecting a known marker. To score transformation events, MEM^R^, CTX^R^ or AMP^R^ colonies were counted. Transformation events were confirmed by measuring the level of resistance to carbapenems (MICs) and other antibiotics to see a change in the resistance phenotype. PCR reactions targeting different resistance genes were also performed (*bla*_KPC_, *strA, strB, aph(3’)*, and *aac(6’)-lb*.

### Antimicrobial susceptibility testing

Disk diffusion assays were used as screening to identify changes in resistance phenotype. To determine the resistance profile of A118, A118::VA360 and A118::Kb18 to ampicillin, amoxicillin/clavulanic acid, cefhalothin, cefoxitin, cefepime, imipenem, MEM, amikacin, gentamicin, nalidixic acid, ciprofloxacin, norfloxacin, tetracycline, and trimethoprim/sulfamethoxazole (Table 1), we followed the Clinical Laboratory Standards Institute (CLSI) guidelines (55). Minimal inhibitory concentrations (MICs) to imipenem, MEM, cefotaxime, ceftazidime, amikacin, gentamicin, kanamycin and tobramycin were determined by the gradient diffusion method (E-test method) (56) with commercial strips (Biomerieux) using the procedures recommended by the supplier (Table 2).

### Whole-genome sequence of A118 clinical strain and A118 transformant cells

Genomic DNA was extracted using a MasterPure DNA Purification kit from Epicentre Biotechnologies. “Shotgun” whole-genome sequencing (WGS) was performed using Illumina MiSeq-I, with Nextera XT libraries for sample preparation. De novo assembly was performed with SPADES assembler version 3.1.0 (57), using a pre-assembly approach with Velvet (58). RAST server was used to predict and annotate open reading frames (59) and BLAST (version 2.0) software was utilized to confirm the predictions. tRNAscan-SE was used to predict tRNA genes (60).

Contig sets of A118, A118::VA360 and A118::Kb18 were ordered and oriented with the Mauve Contig Mover, using the ATCC 17978 genome as reference. Genomes were concatenated clone-wise to generate virtual genomes (61). Sequencing reads were deposited at a local server (http://www.higiene.edu.uy/ddbp/Andres/gtraglia_et_al_2018_data.html)

### Genomic analysis

Sequence analysis was carried out using BLAST (version 2.0) software (http://www.ncbi.nlm.nih.gov/BLAST/). The blast analysis was performed between A118, with A118::VA360, A118::Kb18, A118::Pr and A118::SAC, respectively. The result was sorted by using the R project software, with a 30% minimum identity, 70% minimum coverage and 1e-5 minimum of E-value. The no-codifying sequences inserted into A118 strain were validated with InterProScan. These analyses were done by comparison with protein domains or motifs in the InterProScan database.

The genomic schemes were performed by using Circos and EasyFig softwares (62, 63). ARG-ANNOT, ISfinder and PHAST softwares were used to confirm antibiotic resistance genes, insertion sequences and putative prophage within the sequenced genomes, respectively (64, 65).

BLAST was used to identify resistance genes in *A. baumannii* or *K. pneumoniae* sequences deposited in the GenBank.

### *In-silico* prediction of horizontal gene transfer (HGT) by trees reconciliation

The explicit phylogenetic method was used to analyze potential horizontal genes transfer from one genus to another. Search was based on analysis of topology difference between the phylogenetic trees of gene (protein) clusters and the corresponding phylogenetic species trees (tree reconciliation analysis). To further validate the observation of HGT event detected by explicit phylogenetic methods, we used the information of HGTree database and the NCBI smart blast tool with a parallel BLASTp search to find the closest matches to high-quality sequences. The number of genomes for each analysis were selected per the number of genomes into the HGTree framework available.

### Growth curve of recipient cell and transformant cells

A118, A118::VA360, and A118::Kb18 were grown to stationary phase in LB broth (Fisher BioReagents, Fair Lawn, NJ, USA) with 200 rpm agitation at 37°C. An aliquot (7.5 μl) of A118, A118::VA360 and A118::Kb18 were added to (367.5 μl) LB. Triplicate aliquots (100 μl of this mix were transferred to a 96 well plate. Growth rate curves were generated using a Synergy 2 multi-mode plate reader (BioTek, Winooski, VT, USA) and Gen5 microplate reader software (BioTek), which measured and recorded OD_600_ every 20 minutes. Each condition was tested in triplicates in a non-treated round bottom 96 well polystyrene plate (Costar®, Kennebunk, ME, USA) over 24 hours with light agitation at 37°C. Averages of the triplicates from a single trial were used to report the growth rate curve.

### Quantitative assessment of biofilm formation

A118 and A118::Kb18 strains were grown for 18 hours then diluted 1:100 in LB and 200 μl of these bacterial suspensions were loaded in wells of sterile 96-well polystyrene microtiter plates. LB medium were included as a control. After 24 hours of static incubation at 37°C, the optical density at 595 nm (OD_595_) was measured by using a microplate reader (Multiskan EX, Thermo Electron Corporation, Waltham, MA, USA) and the culture OD_595_ was determined (OD_G_). LB was then removed and each well was washed twice with phosphate buffered saline (PBS). The biofilms were fixed with 100% methanol, stained with 0.5% crystal violet and washed twice with destilled water. After adding 30% glacial acetic acid, the biofilm biomass was measured by reading the OD_595_ (OD_B_). Crystal violet staining levels were expressed relative to the final culture density measured prior to the biofilm assay (biofilm: OD_B_)/OD_G_).

### Protein extraction and separation

*A. baumannii* cultures grown in LB for 18 h were centrifuged at 10,000 × g and 4 °C during 20 min and both pellet and supernatan were saved to extract proteins. To obtain exoproteins, the supernatants were precipitated by adding 10% tricloroacetic acid and incubating at 4 °C for 18 h. The exoproteins were recovered by a centrifuge step at 8,500 × g and 4 °C during 70 min. Then the pellet containing the exoproteins was washed with cold 70% ethanol and resuspended in 0.1 M Tris pH=7.5 and 2 mM PMSF. By the other hand, the pellet obtained at the begining was resuspended in 0.125 M Tris pH=7 and 2% SDS, and heated at 95 °C for 3 min. After centrifuging the suspension at 10,000 × g and 4 °C for 3 min, the protein extracts were recovered from the supernatants. All protein fractions were stored at −80 °C until use. Protein concentration of the samples was determined by Bradford method. After adding Laemmli sample buffer to the samples, they were incubated at 100 °C for 5 min. Then, protein extracts were separated by SDS 10%-PAGE. Polyacrilamide gels were stained with Coomassie blue.

### Killing Assay

A118 and VA360 were used to perform the killing assays. Assays were set up using overnight cultures that were normalized to an OD600 of 1 and done in triplicate. A118 and VA360 were mixed at a predator-prey 1:1 ratio and 5 μl was spotted on a dry LB agar plate. After 4 h, the spot was resuspended in 1 ml saline solution and 10 μl serial dilutions plated on LB agar plate with kanamacyn (20μg/ml). In parallel two other Acinetobacter strains were used (*A. baumannii* A42 strain and *non-baumanniii* A47 strain) (42).

## Acknowledgment

The authors’ work was supported by grants from the Research reported in this publication was supported in part by the National Institutes of Health under awards grants numbers NIH1SC3GM125556-01to MSR; R01AI100560, R01AI063517, R21AI114508, and R01AI072219 to RAB; 2R15AI047115 to MET. This study was supported in part by funds and/or facilities provided by the Cleveland Department of Veterans Affairs, Award Number 1I01BX001974 to RAB from the Biomedical Laboratory Research & Development Service of the VA Office of Research and Development and the Geriatric Research Education and Clinical Center VISN 10 to RAB. The content is solely the responsibility of the authors and does not necessarily represent the official views of the National Institutes of Health or the Department of Veterans Affairs. GMT and CD have a Post-Doctoral Fellowship from CONICET. JF has a SOAR-ELEVAR Scholar Fellowship from Latina/o Graduate Students from the U.S. Department of Education. ASB is a Research Fellow from CONICET. We thank Dr. Luis Actis from the Department of Microbiology, Miami University, Oxford, Ohio, USA and Dr. Fernando Pasteran from the Instituto Nacional de Enfermedades Infecciosas (INEI), ANLIS ‘Dr Carlos G. Malbrán’, Ciudad Autónoma de Buenos Aires, Argentina for providing strains.

**Supplemental material for this article may be found at**

**Figure S1** Bacterial Killing assay. a) Killing assay using VA360 (Prey) and A47, A118, A42 (Predator). Mixtures of VA360/*E. coli* (EcS) kanamycin susceptible were used as control. Representative LB-kanamycin agar plate showing differences in survival of bacterial colonies. b) Killing assay using *E. coli* MG1655-Rif (Prey) and A118 (Predator). MG1655/MG1655-Rif mixtures were used as negative controls. Representative LB-rifampicin agar plate showing differences in survival of bacterial colonies.

**Figure S2** SDS-PAGE of total proteins a) and exoproteins b) of A118 and A118::Kb18 planktonic cultures. The same amount of proteins was loaded in each lane. Gels show three independent experiments (a) or two and three independent experiments (b). MK: molecular weight marker.

**Figure S3** Biofilm formation by A118 and A118::Kb18. Each bar represents the mean ± SD of 6-8 wells from 3 to 4 independent experiments. Biofilm formation values correspond to the OD_595_ of crystal violet (OD_B_) measured relative to the final culture density (OD_G_) after 24 hours.

**Figure S4** Growth rate curves for A118, A118::Kb18, A118::VA360 in LB broth. Optical density (OD_600_) was recorded every 20 minutes for 24 hours and the averages of triplicate samples are shown.

**Table S1** Description of acquired genes identified in *A. baumannii* transformant isolates (A118::Kb18, A118::VA360, A118::Pr and A118::SAC).

**Table S2** Presence of genes in *Klebsiella pneumoniae* sequences deposited in the Genbank database.

**Table S3** Bioinformatic analysis of commonly found *Enterobacteriaceae* or *A. baumannnii* resistance genes.

**Table S4** *Acinetobacter baumannii* isolates harboring genes most generally reported in *Enterobacteriaceae*.

**Table S5** *In-silico* prediction of horizontal gene transfer (HGT) by trees reconciliation data.

